# An evolutionary switch in the specificity of an endosomal CORVET tether underlies formation of regulated secretory vesicles in the ciliate *Tetrahymena thermophila*

**DOI:** 10.1101/210328

**Authors:** Daniela Sparvoli, Elisabeth Richardson, Hiroko Osakada, Xun Lan, Masaaki Iwamoto, Grant R. Bowman, Cassandra Kontur, William A. Bourland, Denis H. Lynn, Jonathan K. Pritchard, Tokuko Haraguchi, Joel B. Dacks, Aaron P. Turkewitz

**Author notes:** joint second authors (Elisabeth Richardson, Hiroko Osakada, Xun Lan). Correspondence should be addressed to: Aaron Turkewitz.

## Abstract

In the endocytic pathway of animals, two related complexes, called CORVET (Class C Core Vacuole/Endosome Transport) and HOPS (Homotypic fusion and protein sorting), act as both tethers and fusion factors for early and late endosomes, respectively. Mutations in CORVET or HOPS lead to trafficking defects and contribute to human disease including immune dysfunction. HOPS and CORVET are conserved throughout eukaryotes but remarkably, in the ciliate *Tetrahymena thermophila,* the HOPS-specific subunits are absent while CORVET-specific subunits have proliferated. VPS8 (Vacuolar Protein Sorting), a CORVET subunit, expanded to 6 paralogs in *Tetrahymena*. This expansion correlated with loss of HOPS within a ciliate subgroup including the Oligohymenophorea, which contains *Tetrahymena*. As uncovered via forward genetics, a single *VPS8* paralog in *Tetrahymena* (*VPS8A*) is required to synthesize prominent secretory granules called mucocysts. More specifically, *∆vps8a* cells fail to deliver a subset of cargo proteins to developing mucocysts, instead accumulating that cargo in vesicles also bearing the mucocyst sorting receptor, Sor4p. Surprisingly, although this transport step relies on CORVET, it does not appear to involve early endosomes. Instead, *Vps8a* associates with the late endosomal/lysosomal marker Rab7, indicating target specificity switching occurred in CORVET subunits during the evolution of ciliates. Mucocysts belong to a markedly diverse and understudied class of protist secretory organelles called extrusomes. Our results underscore that biogenesis of mucocysts depends on endolysosomal trafficking, revealing parallels with invasive organelles in apicomplexan parasites and suggesting that a wide array of secretory adaptations in protists, like in animals, depend on mechanisms related to lysosome biogenesis.

**Abbreviations:**

LRO: (Lysosome-related organelle)
HOPS: (homotypic fusion and protein sorting complex)
CORVET: (Class C core Vacuole/Endosome Transport)
VPS: (vacuolar protein sorting)
GRL: (granule lattice)
GRT: (granule tip)
Igr: (Induced upon granule regeneration)
SNARE: (Soluble NSF attachment protein receptor)
LECA: (last eukaryotic common ancestor)

## Introduction

Cells devote enormous resources to interact with and modify their surroundings. One cellular strategy is to externalize proteins, either by expressing them on the cell surface or by secreting them. Proteins to be secreted are first translocated from the cytoplasm into the endoplasmic reticulum, from whence they are transported through successive membrane-bound compartments, and finally into vesicles[1, 2]. When vesicles fuse with the plasma membrane, called exocytosis, the proteins in the vesicle membrane are exposed on the cell surface while vesicle contents are secreted. In regulated exocytosis, the final exocytic event occurs in response to extracellular stimuli[3, 4].

In animal tissues, multiple classes of vesicles undergo regulated exocytosis to release peptides and other molecules that facilitate fluent cell-cell communication. Dense core granules, such as those in which endocrine hormones are stored for regulated release, arise from the trans-Golgi[5, 6]. A second class of vesicles, which store diverse cargos in different tissues, are called lysosome-related organelles (LROs)[7]. In humans, LROs are vital structures including melanosomes, Weibel-Palade bodies, and T-cell lytic granules[8]. LRO formation depends on trafficking from the trans-Golgi, but LROs simultaneously receive cargo from endosomes[9, 10]. LRO formation involves cytoplasmic and membrane proteins including the small GTPases Rab32/38, SNAREs, the AP3 coat adaptor complex, and a sorting receptor, sortilin/VPS10[11–15]. LRO formation also involves the HOPS complex, a 6-subunit heterohexamer that functions as a multivalent tether between endosomal compartments to facilitate their subsequent fusion[16, 17]. Four HOPS subunits (VPS11, 16, 18, and 33) are also found in a related complex, CORVET, while the remaining 2 subunits in each complex are complex-specific[18, 19]. As shown primarily in budding yeast and animals, CORVET and HOPS are also functionally related, acting as tethers at Rab5- and Rab7-positive endosomes, respectively[20–22]. In mammalian cells, these correspond to successive stages in endosome maturation[23].

Pathways involved in endosomal trafficking and lysosome formation appear to have been present at the time of the last eukaryotic common ancestor (LECA)[24–27]. LECA was a unicellular organism that existed ~1.5 billion years ago, whose membrane compartments have been inferred based on morphological comparisons and genomics-based surveys of compartmental determinants in its descendants, the extant eukaryotes[28] (inter alia). Another inference from such surveys is that many lineages in addition to animals have evolved increasingly complex secretory pathways, but the cell biological details are largely underexplored. Based on microscopy, secretory vesicles in the Alveolate protists, collectively called extrusomes, attracted notice due to their large size, regulated exocytosis, and often elaborate morphologies[29–31]. The Alveolates include largely free-living ciliates and dinoflagellates, and parasitic apicomplexans. Extrusomes in ciliates are functionally and compositionally distinct from those in apicomplexans: the former are used for predation or defense, and perhaps for encystment, while the latter are deployed during host cell invasion[31–36]. However, accumulating evidence indicates that extrusome formation in both ciliates and apicomplexans involves genes associated with LRO biogenesis.

The best-studied apicomplexan extrusomes are the rhoptries in the globally-important parasites *Toxoplasma gondii* and *Plasmodium falciparum[36]*. Rhoptries were initially linked with LROs based on morphological considerations, since they contain internal membranes and thus resemble endosomal multivesicular bodies in animal cells[37]. This idea received molecular support from recent studies in apicomplexans demonstrating involvement of a VPS10/sortilin receptor, early endocytic Rab GTPases, and the VPS11 subunit of the HOPS/CORVET complex[38–41].

Ciliates are an impressively diverse and ecologically important group of microbes, found in nearly all fresh-water, marine and terrestrial environments[42]. The best studied ciliate extrusomes are trichocysts in *Paramecium tetraurelia* and mucocysts in *Tetrahymena thermophila*. The mucocyst lumen is primarily filled with densely-packed Granule lattice (GRL) proteins, which are delivered to immature mucocysts as proproteins and then undergo proteolytic maturation by aspartyl- and cysteine-cathepsins, Cth3p and Cth4p[43, 44]. Processing is essential for secretion, because only processed Grl proteins can form a matrix that expands upon exocytosis to extrude the mucocyst contents. A second family of proteins in mucocysts is called GRT/IGR (Granule tip/Induced on Granule Regeneration)[45, 46].

Significantly, mucocysts are not multivesicular, unlike Apicomplexan rhoptries. However, the delivery to mucocysts of Cth3p and Grt/Igr proteins, but not the Grl proteins, requires the sortilin/VPS10 receptor, Sor4p, as well as an endosomal SNARE (Soluble NSF attachment protein receptor), Stx7l1p (Syntaxin 7-like) [47, 48]. These results suggested that extrusomes in ciliates may depend on LRO-related mechanisms that bear fundamental similarities with those used in apicomplexans. The relationship could be clarified by understanding the role of endosomal tethers in ciliate extrusome formation. Intriguingly, though the HOPS complex is broadly conserved throughout eukaryotes, the HOPS-specific subunits are absent in *Tetrahymena* and were also lost independently in some apicomplexans[49].

One approach to analyzing mucocyst biogenesis is isolating chemically-induced mutants, followed by whole genome sequencing to pinpoint the causative lesions[50]. In this work, we investigate UC616, a mutant previously isolated as part of a large screen[46], and thereby uncover direct evidence for an endosomal intermediate in mucocyst biogenesis. Our data indicate that in *Tetrahymena* a novel CORVET paralog provides HOPS-like function. Our phylogenetic analysis maps the loss of HOPS in *Tetrahymena* and related Oligohymenophorean ciliates, balanced by paralogous expansion in CORVET-specific genes. With these new results, a comparison of regulated secretion in apicomplexans and ciliates indicates that broadly similar mechanisms, evolved in parallel, were employed to generate two dramatically different, highly complex secretory compartments.

## Results

### UC616 bears a recessive mutation that blocks mucocyst formation

The *Tetrahymena* mutant strain UC616 fails to release mucocyst contents upon stimulation (Figure S1A). The defect lies at the level of mucocyst formation, as judged by the failure of proGrl proteins to undergo proteolytic maturation (Figure S1B). In addition, both GRL and GRT proteins were localized aberrantly in UC616 (Figure 1A, 2^nd^ panel). EM analysis revealed small electron dense structures in the cytoplasm of the mutant cells, distinct from wildtype mucocysts (Figure 1B). These structures are similar in appearance (Figure S1C,D) and size (Figure S1E) to Grl3p-containing mucocyst intermediates in the *Δstx7l1* mutant, which is defective in homotypic and/or heterotypic fusion required for mucocyst formation[48]. In standard genetic crosses, all defects in UC616 segregated as expected for a single recessive Mendelian mutation (not shown). Other matings revealed that UC616 is allelic to SB281, a previously-described mutant in mucocyst biogenesis[51, 52], i.e., all UC616xSB281 were fully mutant (not shown).

**Figure 1.**
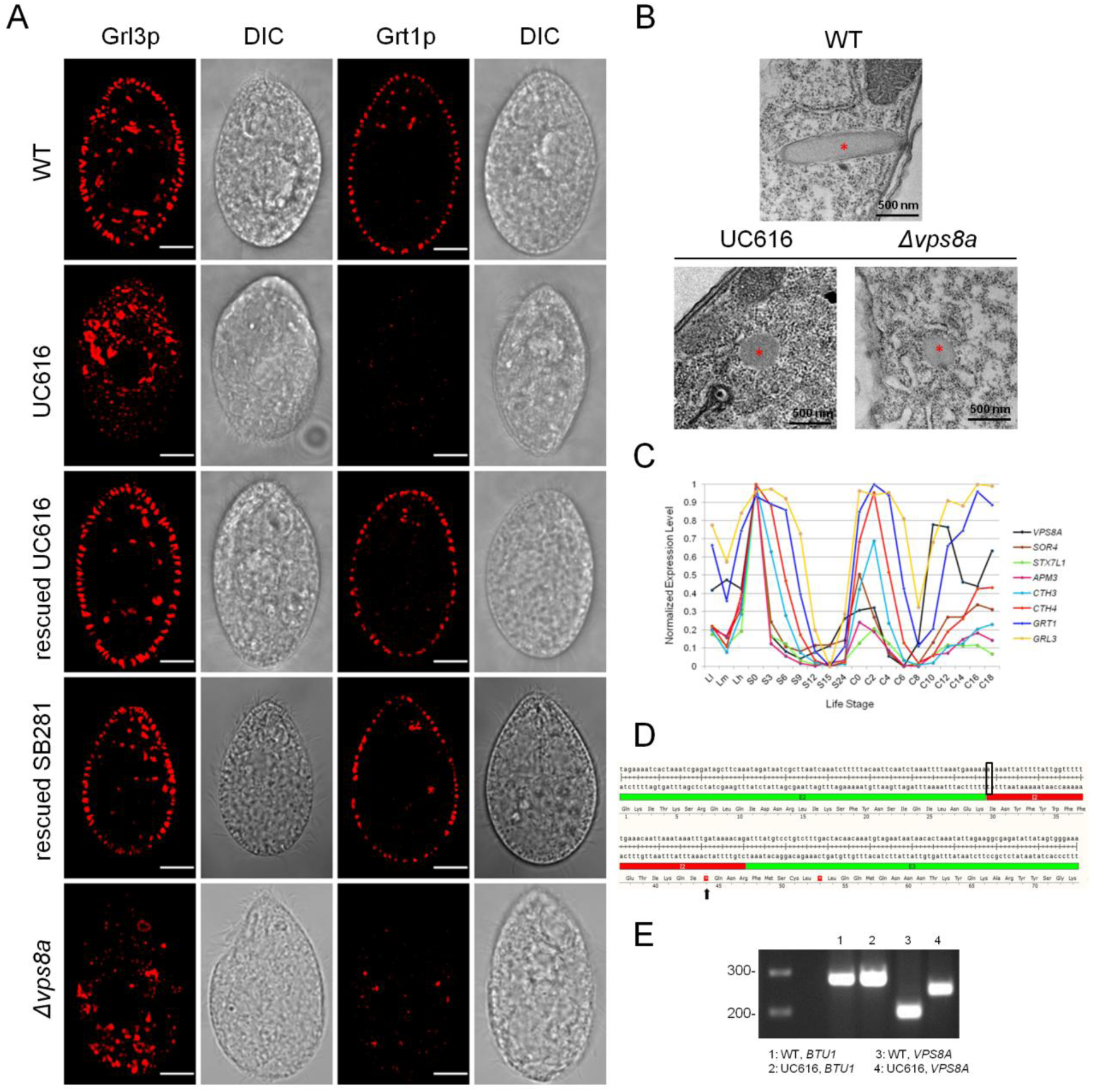
Vps8a is essential for mucocyst biogenesis. A) Confocal cross sections, with paired DIC images, of WT, UC616, rescued UC616 and SB281, and *Δvps8a*. Cells were immunostained with mAbs 5E9 and 4D11, which recognize mucocyst proteins Grl3p (left) and Grt1p (right), respectively. The wildtype pattern of docked mucocysts is absent in UC616 and *Δvps8a*, but was restored in UC616 and SB281 expressing a *VPS8A*-GFP transgene. Scale bars, 10μm. B) Electron microscopy. In WT, mucocysts dock at the plasma membrane. In contrast, UC616 and *Δvps8a* contain cytoplasmic vesicles with granular cores (labeled with asterisks). Scale bars, 500 nm. C) The expression profile of *VPS8A* (black line) is similar to that of mucocyst-related genes *SOR4*, *STX7L1*, *APM3*, *CTH3*, *CTH4*, *GRT1*, and *GRL3*. For the plot, based on data downloaded from tfgd.ihb.ac.cn, each value was normalized to that gene’s maximum expression level. The conditions sampled were growing cultures (low Ll, medium Lm, and high Lh culture density), starvation over 24 hours (S0-S24), and timepoints during conjugation (C0-C18). D) Alignment of the *VPS8A* exon(E2)-intron(I2) junction, and the corresponding translated sequence. The G/A-C/T mutation (black box) results in a premature stop codon (black arrow) 41bp downstream. E) Intron 2 is retained in the *VPS8A* transcript in UC616. Columns 1 and 2 correspond to WT and UC616 cDNA that was PCR-amplified with *BTU1* primers as control; in lanes 3, 4, the primers flanked intron 2 in *VPS8A*. 2% ethidium bromide-stained gel, with 200 and 300 bp DNA fragments indicated to the left. See also Figure S1.

### Sequencing pools of UC616 progeny identifies causative mutation in the *VPS8A* gene

We outcrossed and backcrossed UC616 to obtain progeny, from which we picked single cells that were subsequently expanded to clones. We determined each clone’s phenotype as either mutant or wildtype for regulated exocytosis. Since the mutation is recessive, the mutant allele should be homozygous in the first group of clones but heterozygous or absent in the latter (Figure S1F). Clones from each category were pooled, and their whole genomes sequenced. Single nucleotide variants (SNVs) were then identified that were homozygous only in the mutant pool. Ten such SNVs fell within predicted genes. One of these genes, which we named *VPS8A,* had an expression profile, (downloaded from the Tetrahymena Functional Genomics Database (TFGD, <u>http://tfgd.ihb.ac.cn/</u>)) that is shared with many factors involved in mucocyst biogenesis (Figure 1C)[53]. *VPS8A* is a homolog of pan-eukaryotic VPS8, which encodes one of the two CORVET-specific subunits, i.e., not shared with HOPS[54]. *VPS8A* in UC616 bears a G-to-A mutation predicted to interfere with intron excision (Figure 1D, S1G). We confirmed that intron 576957-577009 (on the reverse strand) is aberrantly retained in UC616 (Figure 1E). This introduces a stop codon 42 nucleotides downstream the splice junction (Figure 1D), resulting in the truncation of the 1499-residue wildtype protein at residue 522, followed by 14 additional novel residues. Surprisingly, genome sequencing of the SB281 mutant identified the identical splice site mutation, while the uniqueness of background SNVs in the two mutant strains confirmed that they are unrelated (not shown).

Both UC616 and SB281 accumulated full cohorts of docked mucocysts after we introduced a wildtype copy of *VPS8A* (Figure S1H; Figure 1A panels 3,4). Such cells were also rescued with regard to proGrl proteolytic processing and exocytosis competence (Figure S1I,J). Moreover, the UC616 phenotypes were reproduced when we disrupted all copies of the Macronuclear *VPS8A* gene (Figure S1K)(Fig. 1A, 5^th^ panel; Fig. S1I,J). *VPS8A* is therefore the locus of the causative mutation in UC616 and SB281, and is required for mucocyst biogenesis. The gene is non-essential and knockout cells (*∆vps8a)* had no growth defect (doubling times: WT 2.6h; *∆vps8a* 2.3h).

### *VPS8A* is required for efficient delivery of a subset of mucocyst proteins

Previous work indicated that pro-Grl proteins appear to be mis-targeted in SB281[46, 52]. We expanded this analysis in the better-defined *∆vps8a* strain, while also taking advantage of recently-described mucocyst components. Specifically, we imaged endogenous Grl3p in pairwise combination with four non-GRL proteins that localize to mucocysts. Three of these (Grt1p, Igr1p, and Cth3p) are soluble proteins in the mucocyst lumen, while the fourth is a membrane-associated SNARE Stx7l1p. GFP-tagged Stx7l1p co-localized equally with Grl3p in both wildtype and mutant cells, indicating that its targeting does not depend on *VPS8A* (Figure S2A,B). In contrast, co-localization of Grt1p and Cth3p (tagged with GFP at the endogenous locus) with endogenous Grl3p were reduced by 78% and 54%, respectively, in *∆vps8a* compared to wildtype (Figure 2A-D).

**Figure 2.**
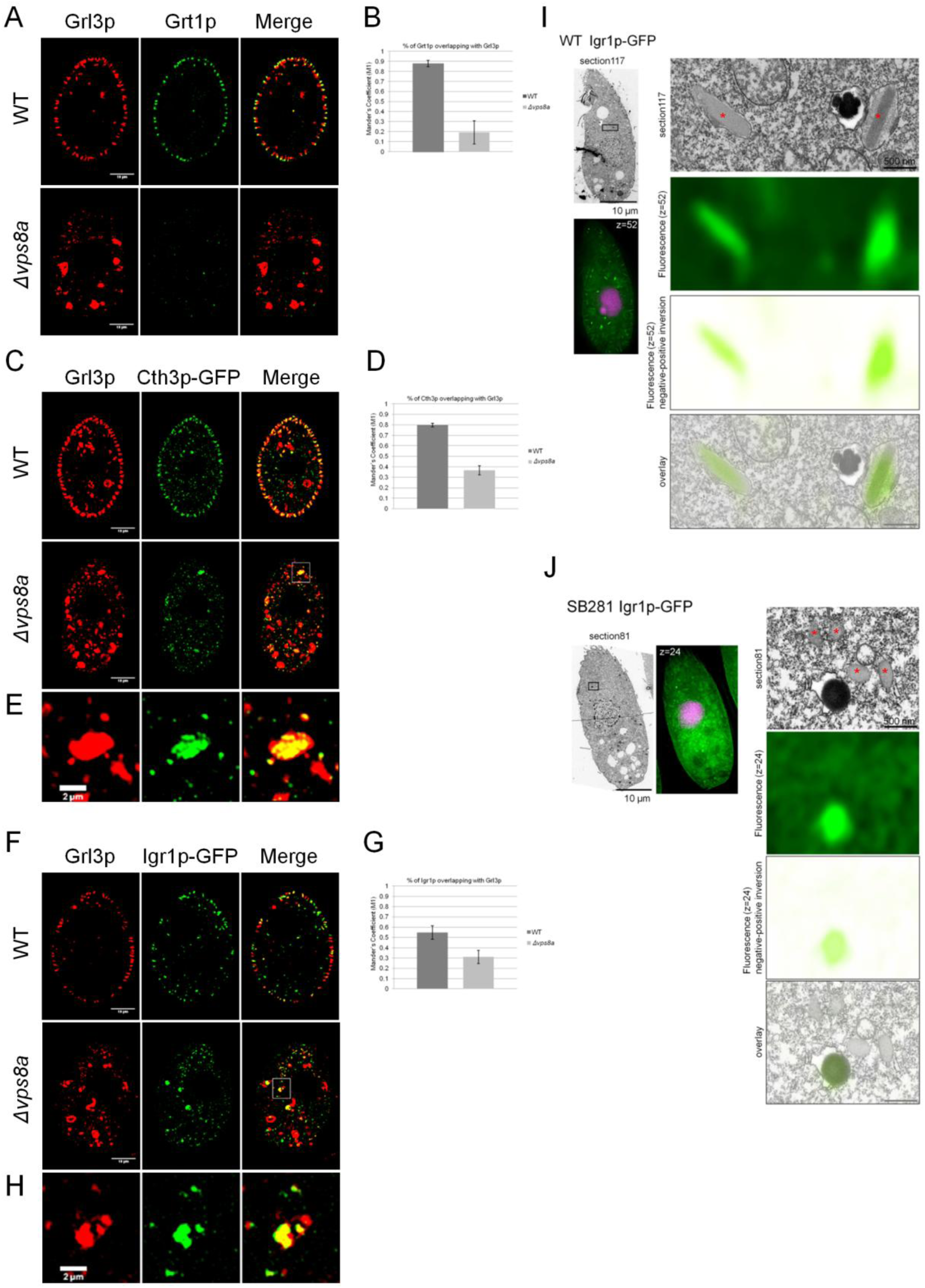
Multiple mucocyst proteins are not delivered to mucocyst intermediates in *Δvps8a* cells. A) WT (upper panel) and *Δvps8a* (bottom panel) cells, co-stained with anti-Grl3p (5E9) and anti-Grt1p (4D11) mAbs conjugated with AlexaFluor 633 and AlexaFluor 488 dyes, respectively. Co-localization between Grt1p and Grl3p is markedly reduced in *Δvps8a*. Confocal cross sections are shown for clarity. Scale bars, 10μm. B) Percentage of overlap between Grt1p and Grl3p (Mander’s coefficient M1), was calculated with 25 non-overlapping images/sample using Fiji-JACoP plugin. C) Localization of Cth3p-GFP in WT and *Δvps8a* cells. Cells were immunolabeled with 5E9 anti-Grl3p and rabbit anti-GFP antibodies. Co-localization of Cth3p-GFP with Grl3p is reduced in *∆vps8a,* and the proteins localize in heterogeneous cytoplasmic vesicles. D) Overlap between Cth3p-GFP and Grl3p was calculated as in (B), based on 40 non-overlapping images/sample. E) Magnification of the boxed area in C (merge), showing co-localization of Cth3p-GFP and Grl3p in a large compartment in *Δvps8a*. F) Localization of Igr1p-GFP in WT and *Δvps8a* cells. Cells were immunolabeled with mAb 5E9. Co-localization of Igr1p-GFP with Grl3p is reduced in *∆vps8a*. G) Overlap between Igr1p-GFP and Grl3p was calculated as in (B), based on 25 non-overlapping images/sample. H) Magnification of the boxed area in F (merge), showing co-localization of Igr1p-GFP and Grl3p in a large compartment in *Δvps8a*. I and J) CLEM imaging of overexpressed Igr1p-GFP in WT and mutant (SB281, which bears the identical *VPS8A* mutation as UC616). Fluorescence images (lower images) are paired with the corresponding electron micrographs (upper images). Left, low magnification images. Scale bar, 10µM. The fuschia-stained structures are the Micro-and Macronuclei stained with DAPI; green represents Igr1p-GFP. Right, boxed regions in the left images are shown at high magnification. From top to bottom: EM image, fluorescence image (Igr1p-GFP),. fluorescence image inverted (negative to positive) and then overlaid on the electron micrograph to show precise mapping of the fluorescence signals onto cellular structures. In WT, Igr1p-GFP is present in mucocysts. In SB281, Igr1p-GFP does not accumulate in the granular mucocyst-related vesicles (asterisks), but is instead found in electron-dense structures likely to represent degradative compartments. Scale bar, 0.5μm. See also Figure S2.

A third lumenal protein, Igr1p (Induced during granule regeneration), was over-expressed as a GFP-tagged copy. Igr1p-GFP showed ~40% reduced co-localization with Grl3p in the mutant compared to wildtype (Figure 2F,G). Significantly, the transport of Grt1p, Igr1p, and Cth3p all depend on Sor4p[47]. Given the known role of CORVET in other organisms in promoting specific endosomal fusion, we therefore hypothesized that Vps8a acts during mucocyst biogenesis to promote heterotypic fusion between a Sor4p-containing compartment and a Grl-containing compartment (Figure 3).

**Figure 3.**
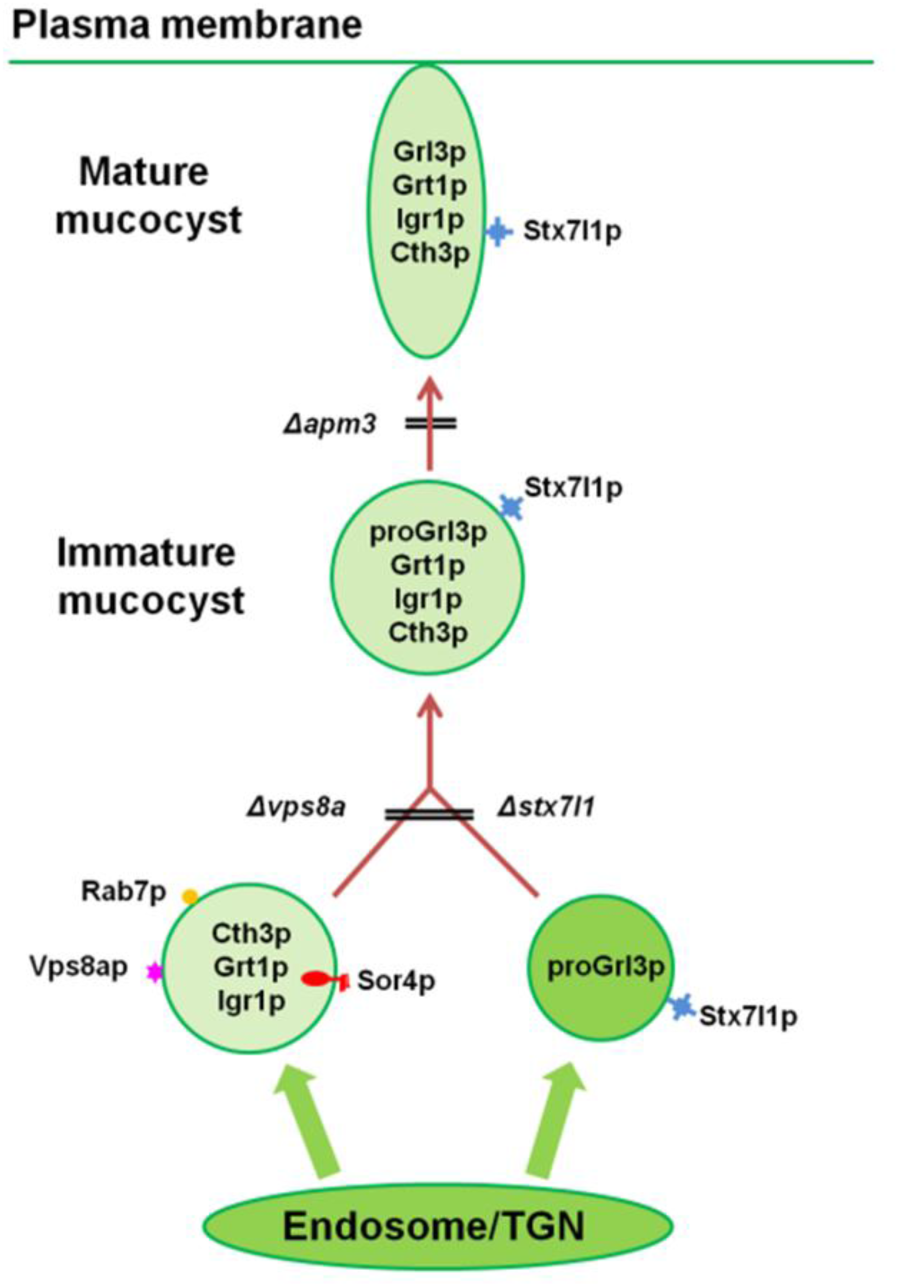
Model for mucocyst biogenesis in *Tetrahymena thermophila*. Two different types of vesicles deliver cargo to immature mucocysts. The first is generated directly from a trans Golgi compartment and contains aggregated proGrl proteins. The second corresponds to a late endosomal compartment, and transports the protease Cth3p and members of the GRT family, Grt1p and Igr1p, bound to the sorting receptor Sor4p. For simplicity, we depict a single compartment that functions both as a trans Golgi and an early endosome, as has been described in plants[80]; whether these are separate compartments in ciliates is unknown. The CORVET subunit Vps8ap (this paper) and the mucocyst-resident syntaxin Stx7l1p mediate the tethering and fusion between the two classes of vesicles. The AP-3 complex appears required for delivery of some as-yet unidentified mucocyst maturation factors.

For Cth3p-GFP and Igr1p-GFP, the co-localization with Grl3p that persists in *∆vps8a* appeared due in large part to inclusion in large structures (Figure 2E,H) distinct from the ~0.4µm spherical vesicles noted earlier (Figure S1C-E). These relatively large compartments are likely to be degradative rather than biosynthetic, based on correlative light and electron microscopy analysis of the Igr1p-GFP expressing cells (Figure 2I,J) and on co-localization with LysoTracker (Figure S2C,D). There was also increased co-localization between Grl3p (expressed as Grl3p-GFP, Figure S2E) with LysoTracker in the *∆vps8a* cells compared to wildtype (Figure S2F,G).

These results suggest that diverse abortive mucocyst intermediates in *∆vps8a* are targeted for degradation.

### *∆vps8a* cells accumulate vesicles containing mis-targeted luminal proteins and their sorting receptor

The mis-targeting of Grt1p, Igr1p and Cth3p in *∆vps8a* can be explained if vesicles containing Sor4p and its bound ligands fail to be delivered to immature mucocysts, so we sought direct evidence for their existence. In wildtype cells, GFP-tagged Sor4p, expressed from the endogenous locus, localized to small mobile cytoplasmic vesicles[47]. Although Sor4p shows robust biochemical interaction with Grt1p and is absolutely required for its delivery to mucocysts[47], the two proteins in wildtype cells show no significant co-localization (Figure 4A,B). In contrast, Sor4p and Grt1p in *∆vps8a* cells co-localize extensively (Figure 4A,B), in vesicles whose size distribution is somewhat shifted compared with Sor4p-containing vesicles in wildtype cells (Figure 4C). The altered localization of Sor4p-GFP in *∆vps8a* does not reflect targeting to degradative compartments, since the puncta do not label with LysoTracker (Figure S3A,B), and Sor4p-GFP in such cells accumulates to higher than wildtype levels (Figure 4D).

**Figure 4.**
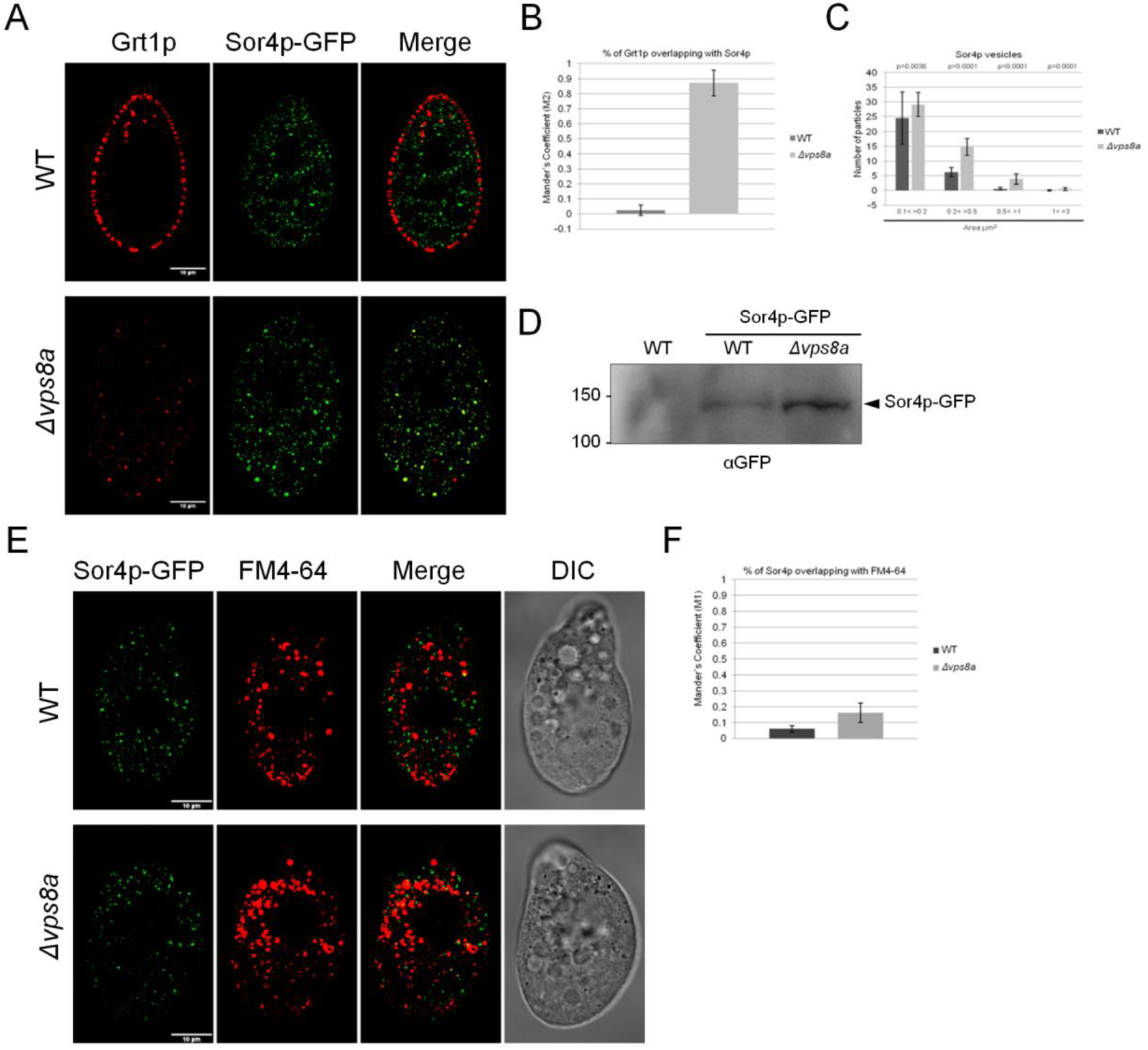
Sor4p co-localizes with its ligand Grt1p in *∆vps8a* but not wildtype cells. A) WT and *Δvps8a* expressing Sor4p-GFP were immunolabeled with anti-Grt1p mAb. Confocal cross sections are shown for clarity. Scale bars, 10μm. B) Overlap between Grt1p and Sor4p-GFP (M2) was calculated as in (2B), based on 40 non-overlapping images/sample. C) The distribution of Sor4p-GFP vesicle sizes is shifted in *∆vps8a* compared to wildtype. Shown are Mean particle counts for each size class, calculated using the Fiji tool “Analyze Particles” with 40 non-overlapping images/sample. p-values determined by two-tailed *t*-test. There are significant increases in vesicle number in *∆vps8a* compared to WT in multiple size classes: (0.1< >0.2 μm^2^) = 0.6-fold; (0.2< >0.5 μm^2^) = 1.6-fold; (0.5< >1 μm^2^) = 5.8-fold. In addition, rare Sor4p-GFP-positive structures of (1< >3 μm^2^) appear almost exclusively in *∆vps8a*. D) SDS-PAGE and Western blotting of whole cell lysates from 1.5x10^5^ WT and *Δvps8a* cells expressing Sor4p-GFP, and an untransformed WT control, probed with anti-GFP antibody. The position predicted for full-length Sor4-GFP is indicated. *∆vps8a* cells accumulate more Sor4p-GFP than do WT cells. E) Confocal live microscopy of WT and *Δvps8a* expressing Sor4p-GFP and pulse labeled with FM4-64 for 5min. A small fraction of Sor4p-GFP vesicles in WT cells is labeled with FM4-64, and this overlap increases in *Δvps8a*. Scale bars, 10 μm. F) Overlap of Sor4p-GFP with FM4-64 (M1) was calculated as in Figure 2B, using 40 non-overlapping images/sample. See also Figure S3.

These puncta may represent transport intermediates that are transient in wildtype cells, but trapped in *∆vps8a* because they are blocked in fusion with their acceptor compartment. They appear to derive from endosomes, as suggested by the ~ 2.7-fold and ~ 5.1-fold increased co-localization of the endocytic tracer FM4-64 with Sor4p-GFP or Igr1p-GFP, respectively, in

*∆vps8a* compared to wildtype (Figure 4E,F; S3C,D). Importantly, in the *∆vps8a* cells, Sor4p-GFP shows no significant co-localization with Grl3p (Figure S3E,F), consistent with the idea that Grl proteins are delivered to mucocysts via a different route from Sor4p cargo. Taken together, these results support the model in Figure 3.

### *VPS8* underwent paralogous expansion and functional diversification in the ciliate lineage containing *T. thermophila*

VPS8 is represented by a single gene in *H. sapiens*, *S. cerevisiae, D. melanogaster*, *T. gondii*, and many other eukaryotes. In contrast, six *VPS8* paralogs exist in *T. thermophila*. Our phylogenetic analysis showed that at least four paralogs were maintained since the ancestor of the Oligohymenophorean ciliate lineage (Figure S4A). The root of the tree was determined using *Oxytricha trifallax* Vps8 sequences as an outgroup, and the monophyly of the Oligohymenophorea-containing clade to the exclusion of Oxytricha had complete support under both MrBayes and RAxML (1/100). This analysis indicates that the post-Spirotrichea expansions occurred independently of expansions in other lineages, and led us to postulate distinct functional roles for individual VPS8 paralogs that arose in Oligomenophorea. Consistent with this idea, the six *T. thermophila* paralogs have non-identical expression profiles (Figure 5A), and the *VPS8A* profile appears unique. To test whether functional diversification had occurred between paralogs, we targeted each of the additional five paralogs for knockout in the Macronucleus. For two of the genes, we could not recover viable knockout cells, i.e., a drug resistance cassette integrated at the Macronuclear locus could not be driven to fixation (Figure S4B). These results indicate that two paralogs are essential, unlike *VPS8A*. Targeted disruption of the remaining three paralogs revealed that complete loss of *VPS8B*, *E* or *F* had no effect on mucocyst secretion (Figure 5B, S4C). Thus, *VPS8A* represents a CORVET subunit that appears specialized for a single pathway of endolysosomal trafficking, and may have little functional overlap with other *VPS8* paralogs.

**Figure 5.**
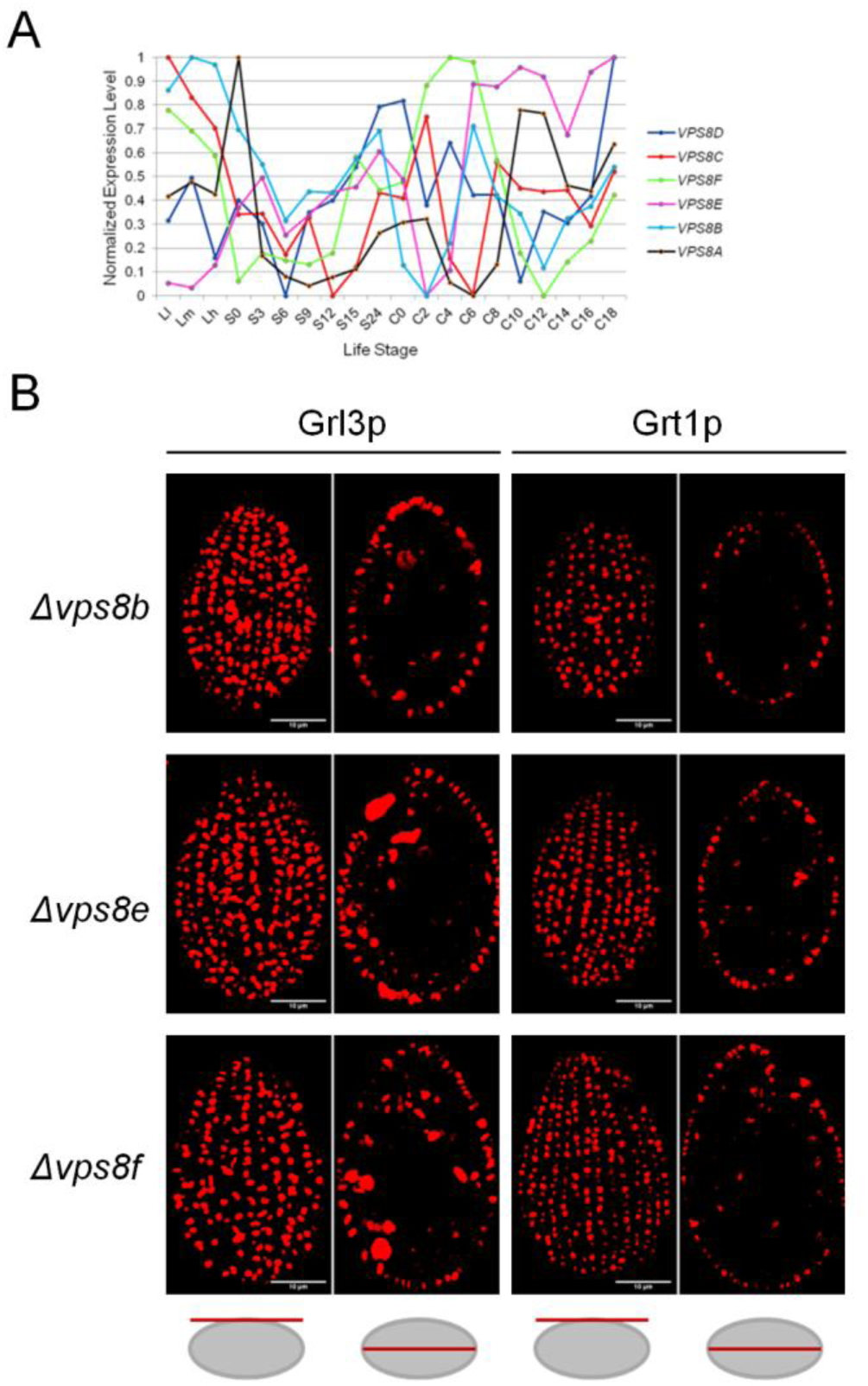
Expansion and diversification in ciliates of CORVET *VPS8* genes. A) Expression profiles of the six *T. thermophila VPS8* paralogs, displayed as in (1C), shows that each paralog has distinct expression peaks, and only *VPS8A* (black line) shows the profile associated with mucocyst formation (Figure 1C). Expression data were downloaded from the Tetrahymena Functional Genomics Database (TFGD, <u>http://tfgd.ihb.ac.cn/</u>). B) Non-essential paralogs *VPSB*, *E*, and *F* are dispensable for mucocyst formation. *Δvps8B, Δvps8E*, and *Δvps8F* cells were immunolabeled with antibodies against Grl3p or Grt1p, followed by Texas red-conjugated goat anti-mouse. All strains maintained WT patterns of docked mucocysts. Surface and cross sections are indicated at the panel bottom. Scale bars, 10μm. See also Figure S4.

As tethers, CORVET complexes bridge two membranes by interacting with RabGTPases and SNAREs on the target organelles[55–57]. In fungi and animals, the Vps8 subunit makes direct contact with Rab GTPase in a target membrane[54]. Thus, the six *VPS8* paralogs in *Tetrahymena* may confer different target specificities for distinct CORVET complexes. Moreover, three of the four core subunits found in both CORVET and HOPS, though not the CORVET-specific subunit *VPS3*, are also present as multiple paralogs in *T. thermophila*: *VPS16*, *18*, and *33*[49]. To ask whether Vps8ap-containing complexes are enriched in specific paralogs of other subunits, we performed pulldowns from cells transformed to express Vps8a-ZZ-FLAG in combination with an HA-tagged copy of either Vps16a or b. Our results indicate that Vps8ap preferentially associates with Vps16bp (Figure S4D,E), suggesting that different CORVET complexes contain distinct paralogous subunits. Hence different CORVET paralogs may, in a combinatorial fashion, partition endosomal traffic between a variety of pathways.

### Vps8a is associated with Rab7-positive endosomes

If the Sor4p-Grt1p-positive vesicles in *∆vps8a* cells are related to early endosomes, we would expect them to label readily with an endosomal tracer. However, they showed only limited overlap with FM4-64 (Figure 4E,F). We considered this result in light of the fact that *T. thermophila* lacks all subunits specific for the HOPS complex. By analyzing genomes and transcriptomes from across the diversity of ciliates, we found that loss of HOPS occurred after divergence of Spirotrichea from the subsequent lineages including Oligohymenophora, the group in which the CORVET paralogous expansion took place (Figure 6A). This raised the possibility that novel CORVET complexes were providing HOPS-like functions, and acting at late endosomes. To test this idea, we looked at co-localization with RabGTPases.

**Figure 6.**
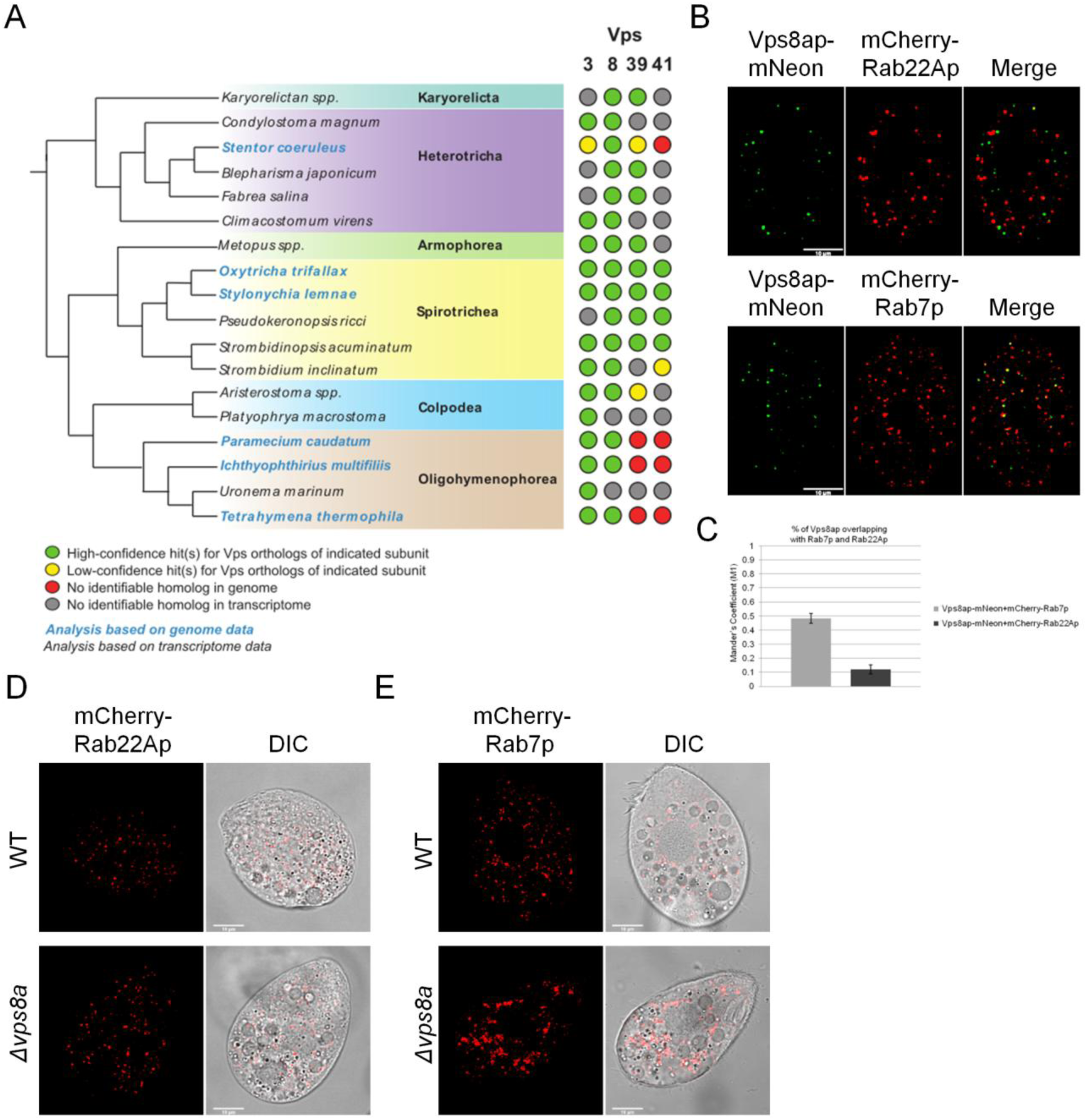
Association of Vps8ap with Rab7 endosomes. A) Distribution of HOPS/CORVET subunits across ciliate diversity. Relationships between ciliate species, based upon [81], are depicted by the evolutionary tree on the left (not to scale), with phylogenetic class indicated in bold. The dotplot on the right depicts the presence/absence of the indicated subunit within each species, as defined in Experimental Procedures. Gene IDs are listed in Table S1. B) Cells were transformed to co-express mNeon-tagged Vps8ap at the endogenous locus together with either the Rab5 homolog mCherry-Rab22Ap (upper panel) or mCherry-Rab7p (lower panel), respectively. Rab transgene overexpression was induced with 1 μg/ml CdCl_2_ for 2.5h in SPP. Scale bars, 10μm. C) The overlap between Vps8ap-mNeon and each Rab-GTPase (M1) was measured as in Figure 2B. Vps8a-mNeon overlapped ~50% with mCherry-Rab7, but only ~12% with mCherry-Rab22A. The Mean M1 values were derived from 65 and 66 non-overlapping images for Rab22Ap and Rab7p samples, respectively. D) and E) Endosome distribution in WT vs *∆vps8a* cells. Shown are single frames from time-lapse movies of WT and *Δvps8a* cells, overexpressing mCherry-Rab22Ap (D) and mCherry-Rab7p (E). Cells were transferred to S-medium and mCherry-Rab transgene overexpression was induced as in (B). Rab22Ap-positive endosomes have a similar distribution in WT and *∆vps8a*, but Rab7p-positive endosomes form large clusters in *∆vps8a* cells. Scale bars, 10μm. See also Figure S5.

In animal cells, CORVET is associated with Rab5-positive early endosomes, which mature to Rab7-and HOPS-associated late endosomes[20]. CORVET also associates with Rab5 in the fungi *S. cerevisiae* and *Aspergillus nidulans*, which in the latter marks early endosomes[21, 58]. Rab22 is the Rab paralog most closely related to Rab5 [59]. *T. thermophila* lacks a clear Rab5 ortholog, but Rab22A co-localizes extensively with the bulk endosomal tracer FM4-64, consistent with it marking early endosomes, while the sole Rab7 homolog associates with vesicles that accumulate LysoTracker and likely represent late endosomes/lysosomes[60].

We therefore analyzed co-localization of mNeon-tagged Vps8a with mCherry-tagged Rab22A or Rab7. Vps8a-mNeon was integrated to fully replace *VPS8A* at the endogenous locus. The tagged protein was functional, judging by the fact that the cells accumulated docked mucocysts, and mucocyst secretion was nearly equivalent to wildtype (Figure S5A-C). Strikingly, Vps8a showed roughly 50% co-localization with Rab7, but only ~10% co-localization with Rab22A (Figure 6B,C). These data, combined with our results on FM4-64 accumulation, strongly suggest that the Vps8a-containing CORVET complex in *T. thermophila* acts primarily at a late endosomal compartment.

We then asked whether the absence of Vps8ap led to a change in the Rab22A or Rab7 compartments compared to wildtype. To this end, we characterized the size and number of Rab22A and Rab7 puncta in both live and fixed wildtype and *Δvps8a* cells, and measured particle number and size distributions in Z-stacks of the fixed samples. The Rab22A-positive puncta had a similar distribution and appearance in the mutant and wildtype (Figure 6D; S5D). Vesicles in the medium and larger size classes were more numerous in the mutant (Figure S5E). Rab7-positive structures showed a more striking difference in the wildtype vs *∆vps8a*. In wildtype, Rab7 puncta are dispersed in the cytoplasm, but in *∆vps8a* they form large clusters (Figure 6E; S5F). Consistent with this, we measured a mild increase in the number of Rab7 small, medium, and large puncta in *∆vps8a*, but a significantly larger increase in a size class that corresponds to clusters (Figure S5G). This was also seen in live cells (not shown).

## Discussion

Guided by results of a forward genetic screen, we discovered that mucocyst formation in *T. thermophila* requires the *VPS8A* gene, a subunit of the CORVET complex. The *∆vps8a* cells accumulate vesicles that contain the mucocyst sorting receptor Sor4p as well as several cargo proteins including Grt1p, which in wildtype cells are delivered to immature mucocysts. These vesicles are likely to represent transient intermediates in mucocyst biogenesis that accumulate in the mutant because tethering and fusion to their target is inhibited. Thus, we hypothesize that the primary defect in ∆*vps8a* is in the fate of endosome-related vesicles that deliver proteins including maturation factors to immature mucocysts. The involvement of CORVET in mucocyst formation reinforces and extends the prior evidence that this pathway relies on endosomal trafficking[47, 48]. Our results support a model of parallel sorting/delivery of mucocyst contents: GRL proteins are segregated and sorted from other secretory cargo based on aggregation, while GRT/IGR proteins, as well as the processing cathepsin Cth3p, are delivered via endosomes as ligands of the Sor4p receptor.

An additional defect was previously documented in the SB281 strain, which we discovered also bears a mutation in *VPS8A*. Starved SB281 cells show constitutive secretion of pro-GRL proteins that in wildtype cells are instead efficiently stored and then proteolytically processed[52]. Interestingly, perturbing the maturation of regulated secretory compartments in animal cells also leads to aberrant constitutive secretion [61]. In *Tetrahymena*, a fraction of the mucocyst cargo proteins in the mutant cells is also found in degradative compartments, which could reflect autophagy of abortive mucocyst intermediates.

CORVET as well as HOPS genes are found throughout eukaryotes[25], implying that both complexes arose during pre-LECA evolution, with duplication and co-evolution of genes producing two complexes whose subtly-altered specificities contributed to differentiation of endosomal trafficking pathways. Prior to this paper, functional characterization of CORVET has only been reported in a single eukaryotic lineage, the Opisthokonts (including animals and fungi), where it associates with Rab5/VPS21-positive endosomes. HOPS has recently been characterized in a second lineage, plants (lineage Archaeplastida), where it appears to function at late endosomes/vacuoles[62, 63]. We previously noted the unusual absence of HOPS-specific subunits in genomes of two ciliates, *T. thermophila* and *Ichthyophthirius multifiliis*[49]. By analyzing additional ciliate genomes, we can now infer that those losses occurred prior to the expansion of the Oligohymenophorea. These losses are unlikely to reflect an evolutionary reduction in the complexity of endosomal pathways: *T. thermophila* expresses at least three Rab8 paralogs and six likely Rab11 paralogs, and HOPS-specific losses are paralleled by proliferation of both core and CORVET-specific subunits. The six *VPS8* paralogs may be specialized for distinct endocytic pathways, reflected by unique peaks within each expression profile. We found that only the *VPS8A* paralog is essential for mucocyst formation, while no other non-essential paralogs showed mucocyst phenotypes. Moreover, two of the *VPS8* paralogs are essential, while the others have no associated growth phenotypes.

The function of HOPS/CORVET depends on each complex bridging two different vesicles, by making multivalent contacts with vesicle constituents including Rabs, SNAREs, coat proteins, and phospholipids[64, 65]. In the prevailing model for yeast CORVET, the recognition of Rabs on the two target membranes occurs via the Vps8 and Vps3 subunits[66–71]. Thus, in ciliates the expansion of VPS8 paralogs may have contributed to specifying a large set of endolysosomal compartments, if different VPS8 paralogs preferentially bind to different Rabs. Consistent with this idea, RabGTPases also underwent paralogous expansions in Oligohymenophorea[60, 72-74]. Intriguingly, the 2^nd^ well-established Rab-binding subunit in the yeast model, Vps3, is encoded by a single gene in *T. thermophila*, suggesting that additional specificity may be determined by interactions with other vesicle proteins, e.g., SNAREs that also expanded in Oligohymenophorea. In yeast, both Vps16 and Vps33 are involved in binding to SNAREs [75], and both genes are encoded by multiple paralogs in *T. thermophila*. While we have not yet physically characterized a *T. thermophila* CORVET complex, we found that Vps8a could be robustly co-precipitated with the core subunit Vps16b, consistent with the idea that the subunits assemble as they do in other studied organisms. It is worth noting, however, that functional CORVET sub-complexes have been reported in Drosophila [76]. How many distinct CORVET complexes exist in *Tetrahymena*? In addition to six *VSP8* paralogs, *Tetrahymena* also expresses two paralogs each of the core subunits *VPS16* and *33*, and four paralogs of core subunit *VPS18*, so in theory could generate 96 compositionally-distinct CORVET complexes.

A key finding was that Vps8a co-localizes with the late endosomal Rab7 rather than with the early endosomal Rab22. The basis for the change in *VPS8* specificity, e.g., whether Vps8a directly interacts with Rab7, remains to be determined. Given that loss of HOPS occurred after the Oligohymenophorea/Spirotrichea split but before the expansion of Oligohymenophorea, an attractive hypothesis is that new CORVET paralogs in this lineage took on HOPS-like specificity. The precise role of Vps8a in *Tetrahymena* is not yet clear. The *∆vps8a* cells show marked clustering of Rab7-positive vesicles. Interestingly, these are unlikely to be strongly acidic since they do not label with LysoTracker; i.e., there is no comparable clustering of LysoTracker-positive vesicles in the mutant cells. Surprisingly, Vps8a did not appreciably co-localize with Grl3p in immature or mature mucocysts. The absence of Vps8a at the presumed acceptor compartment may indicate that the tethering step *per se* is very short lived. However, another possibility is that the Sor4p-GFP vesicles bearing mucocyst cargo first undergo Vps8a-dependent fusion with an unknown compartment, prior to cargo delivery at immature mucocysts.

In animals, mechanisms involved in lysosome biogenesis are deployed to generate a remarkably diverse collection of complex secretory organelles including melanosomes, pigment granules, Weibel-Palade bodies, and T-cell lytic granules[8]. In the apicomplexan *Toxoplasma gondii*, formation of three types of secretory organelles including rhoptries was inhibited by knockdown of VPS11, a core subunit of both HOPS and CORVET[40]. Although the authors assumed that CORVET is absent in *T. gondii*, our previous analysis of HOPS and CORVET subunit evolution in apicomplexans would argue that phenotypes in *Toxoplasma* VPS11 knockdown cells could reflect the loss or modified function of either or both complexes[25]. Most importantly, the requirement for CORVET and/or HOPS underscores the relevance of LRO mechanisms in apicomplexan regulated secretion. Moreover, regulated secretion in dinoflagellates is also likely to share a common origin with ciliates[77].

*Tetrahymena* mucocysts represent an elaborate secretory compartment, generated via a complex stepwise program. GRL proteins assemble in the mucocyst lumen to form an elongated crystal, which expands in a spring-like fashion during exocytosis when exposed to extracellular calcium[78]. Importantly, assembly occurs only after the pro-GRL proteins undergo processing, and the key processing enzyme, Cth3p, is delivered to mucocysts via the *SOR4*/*STX7L1*/*VPS8A*-dependent pathway[47, 48, 79]. In these cells, mechanisms of LRO biogenesis are thus used to process secretory cargo using lysosome-related hydrolases to assemble calcium-sensitive springs. Our results, taken together with findings in apicomplexans, indicate that not only animals, but also protists in the distantly related and globally important Alveolate lineage, adapted LRO-based strategies to enhance their secretory responses.

## Experimental Procedures

### Cell strains and culture conditions

*Tetrahymena thermophila* strains used in this work are indicated in Supplemental Table S3. Cells were grown overnight in SPP (2% proteose peptone, 0.1% yeast extract, 0.2% dextrose, 0.003% ferric-EDTA) supplemented with 250 ug/ml penicillin G, 250 ug/ml streptomycin sulfate, and 0.25 μg/ml amphotericin B fungizone, to medium density (1-3 × 10^5^ cells/ml). For biolistic transformation, growing cells were subsequently starved in 10 mM Tris buffer, pH 7.4. Fed and starved cells were both kept at 30 °C with agitation at 99 rpm. For live microscopy, grown cells were transferred to S medium (0.2% yeast extract, 0.003% ferric-EDTA) for 2-4 hours prior to imaging. Culture densities were measured using a Z1 Coulter Counter (Beckman Coulter Inc.). **Genetic methods**

The methods used for genetic crosses, selecting progeny, and mating type determination are as previously detailed[50].

### Testing assortants for exocytosis competence

Assortants were tested for the ability to secrete mucocysts in response to the secretagogue Alcian blue, using a 96-well plate format and subsequently in bulk cultures, as described previously[50].

### Genomic DNA extraction

Genomic DNA was extracted as previously described[50].

### Whole genome sequencing

The genome sequencing of the F2 pools was performed as previously described[50]. A total number of 295 million paired-end reads (2 × 100 bp) were generated. The F2 lines with the mutation (UC616) of interest were sequenced to ~147-fold genome coverage. The F2 lines showed the wildtype phenotype (WT) were sequenced to ~180-fold genome coverage. Strain SB281 was sequenced to ~198-fold genome coverage. Summary of UC616 and SB281 genome DNA sequencing data is shown in Supplemental Table S4. Genome sequencing of *Tetrahymena* primarily reflects the Macronuclear genome in which genes are generally present at ~45 copies, compared to the two copies in the Micronucleus. Because the Micronucleus represents the germline nucleus, only Micronuclear alleles are transmitted to progeny[82].

However, because all parental lines are wildtype for exocytosis except for UC616 itself, we made the simplifying assumption that bulk DNA sequencing of phenotypically mutant vs. wildtype F2 lines would identify the UC616 causative mutation.

### Sequence alignment

For the analysis of the sequenced genomes, reads from total genomic DNA sequencing were mapped to the *T. thermophila* Macronuclear genome sequence released by Broad Institute using the Burrows-Wheeler Aligner software (BWA) version 0.7.10 [83–85]. Default parameters were used when running bwa mem except the following: 1) Adding proper read group IDs for each sample; 2) Enabling “Mark shorter split hits as secondary” using the – M option. The output files were sorted by coordinates and converted to bam files using the Picard Tools version 1.111. Picard Tools was also used to remove PCR duplicates and index the bam files. To align the *VPS8A* genomic region (TTHERM_00290710, from TGD, <u>http://ciliate.org</u>), containing the identified SNV, and the corresponding transcript (TranscriptID_000003408, from TFGD, <u>http://tfgd.ihb.ac.cn/</u>), we used **MU**ltiple **S**equence **C**omparison by **L**og-**E**xpectation tool (MUSCLE) version 3.8.31.

### Variant discovery

The Genome Analysis Toolkit (GATK) version 3.3.0 was applied to identify variants with total genomic DNA sequencing data of the three samples [86]. The following steps were taken in this procedure. 1) Realignment of reads using “RealignerTargetCreator” and “IndelRealigner” tool to correct the misalignment caused by site mutations, insertions and deletions; 2) haplotypes of each sample were called using the “HaplotypeCaller” tool with the “emitRefConfidence” set to “GVCF”, “variant_index_type” set to “LINEAR”, “variant_index_parameter” set to “128000”, “genotyping_mode” set to “DISCOVERY”, “stand_emit_conf” set to “10”, and “stand_call_conf” set to “30”; and 3) variances in the three samples were jointly called using the “GenotypeGVCFs” tool. Full documentation of parameters can be found at <u>https://www.broadinstitute.org/gatk/guide/</u>tooldocs/org_broadinstitute_gatk_tools_walkers_haplo typecaller_HaplotypeCaller.php#-output_mode. A total number of 51,121 variants was called with this procedure. Among these, 31,744 were Single Nucleotide Variants (SNVs), which resulted in an SNV density of ~0.31 kbp^−1^ for a genome of a mappable size of 103,014,375 bp[83].

### Candidate screening

The candidate causal variants of the exocytosis deficient phenotype were selected as previously described[50]. A total of 35 such candidate sites was found in UC616. Among these 35 SNVs, 10 lay within coding regions or at intron-exon junctions.

### Transcription profiles

Gene expression profiles were downloaded from the *Tetrahymena* Functional Genomics Database (TFGD, <u>http://tfgd.ihb.ac.cn/</u>)[87, 88]. For plotting the graphs, each profile was normalized by setting the gene’s maximum expression level to 1.

### Generation of *VPS8* knockout strains

The Macronuclear open reading frame (ORF) of each of the *VPS8* paralogs (TTHERM_00290710, *VPS8A*; TTHERM_00393150, *VPS8B*; TTHERM_00716100, *VPS8C*; TTHERM_00532700, *VPS8D*; TTHERM_00691590, *VPS8E*; TTHERM_00384890, *VPS8F*) was replaced with the paromomycin (Neo4) drug resistance cassette [89] via homologous recombination with the linearized vectors pVPS8AMACKO-Neo4, pVPS8BMACKO-Neo4, pVPS8CMACKO-Neo4, pVPS8DMACKO-Neo4, pVPS8EMACKO-Neo4, pVPS8FMACKO-Neo4. PCR was used to amplify 500-800bp of the genomic regions upstream (5’UTR) and downstream (3’UTR) of each ORF. The amplified fragments were subsequently cloned into specific restriction sites, flanking the Neo4 cassette of the pNeo4 vector, by Quick Ligation (New England, Biolabs Inc.) unless otherwise specified: NheI/PstI and EcoRV sites for TTHERM_00290710 [the 3’UTR was introduced in the linearized pNeo4 vector by In-Fusion cloning kit (Clontech, Mountain View, CA), at the EcoRV site]; SacI/PstI and XhoI/ApaI sites for TTHERM_00393150, TTHERM_00716100, TTHERM_00532700; NheI/PstI and XhoI/EcoRV sites for TTHERM_00691590, TTHERM_00384890. The primers used to create these constructs are listed in Supplemental Table S5. Each construct was linearized by digestion with SacI and KpnI and transformed into CU428.1 cells by biolistic transformation.

### RT-PCR assessment of *VPS8* disruption, and intron retention in the *VPS8A* transcript of UC616

3x10^5^ cells from overnight cultures were pelleted, washed once with 10mM Tris, pH 7.4, and total RNA was isolated using RNeasy Mini Kit (Qiagen, Valencia, CA) according to the manufacturer’s instructions. The cDNA synthesis, from 2 μg of total RNA, was performed as per manufacturer’s instructions using High-Capacity cDNA Reverse Transcription Kit (Applied Biosystems, Foster City, CA). The cDNA was PCR amplified to assay the presence of the corresponding transcripts (200-250bp) in the knockout strains, and the retention of the intron in the *VPS8A* transcript (250bp) of UC616 and SB281, using primers listed in Supplemental Table S5. To confirm that equal amounts of cDNA were being amplified, reactions with primers specific for β-tubulin 1 (*BTU1*) were run in parallel.

### Expression of Vps8ap-GFP in mutant strains

*VPS8A* was C-terminally fused to monomeric enhanced GFP (mEGFP), and reintroduced at the endogenous *VPS8A* locus in UC616 and SB281 by homologous recombination, using pVPS8A-mEGFP-Neo4. To rescue the *VPS8A* mutation in UC616 and SB281 by expressing a wildtype copy of the gene, 4352bp (of a total of 5091bp) of the *VPS8A* genomic sequence (minus the stop codon), were cloned in the linearized pmEGFP-Neo4 vector [47] at the BamH1 site, by In-Fusion cloning (Clontech, Mountain View, CA). The construct also contains 565bp of *VPS8A* downstream genomic sequence, following the Neo4 drug resistance cassette, cloned in the linearized vector at the EcoRV site, as described earlier for the pVPS8AMACKO-Neo4 vector. The construct was then linearized with NotI and KpnI, and introduced in the Macronucleus by biolistic transformation. The two *VPS8A* fragments were amplified by PCR with primers listed in the Supplemental Table S5.

### Expression of Grl3p-GFP, Cth3p-GFP and Sor4p-GFP at the corresponding endogenous locus in *Δvps8a*

The pSOR4-mEGFP-Neo4 vector [47] was used as template to create pSOR4-mEGFP-Chx and pCTH3-mEGFP-Chx vectors. The former was obtained by replacing the Neo4 cassette in the pSOR4-mEGFP-Neo4 vector, with the cycloheximide (Chx) drug resistance cassette [90]. The Chx cassette was sub-cloned in the PstI and EcorV sites of the digested pSOR4-mEGFP vector. The pCTH3-mEGFP-Chx vector was created by replacing the 688bp and 625bp *SOR4* genomic fragments in the pSOR4-mEGFP-Chx construct, with the corresponding 817bp (minus the stop codon) and 710bp *CTH3* fragments, following digestion with SacI/BamHI and EcoRV/XhoI, respectively. All fragments were PCR amplified using primers listed in Supplemental Table S5, and the sub-cloning was performed by Quick Ligation (New England, Biolabs Inc.). The pSOR4-mEGFP-Chx, pCTH3-mEGFP-Chx, and the previously-described pGRL3-smGFP-2myc-6His-Chx vectors[48], were linearized with SacI and KpnI, and transformed in *Δvps8a* by biolistic transformation.

### Endogenous tagging of *VPS8A* with mNeon

Two mNeonGreen fluorescent tags were fused at the C-terminus of the *VPS8A* macronuclear ORF via homologous recombination using linearized pVPS8A-2mNeon-6myc-Neo4. First, the p2mNeon-6myc-Neo4 vector was constructed. The 2xmNeon tag, followed by 6xc-myc tag, was PCR-amplified from pFAP256-mNeonGreen-6myc-BirA vector (a gift from J. Gaertig, University of Georgia, Athens, GA) and introduced at the SpeI site of a modified version of pNeo4, containing the *BTU1* terminator upstream of the Neo4 cassette, by In-Fusion cloning (Clontech, Mountain View, CA). The C-terminal 828 bp of the *VPS8A* genomic locus (lacking the stop codon) and, subsequently, the 565 bp-long downstream genomic region of *VPS8A*, were cloned at the SacI/MluI and EcoRV sites of the digested p2mNeon-6myc-Neo4 vector, respectively.

The final vector was digested with SacI and KpnI prior to biolistic transformation of CU428.1. The primers are listed in Supplemental Table S5.

### Endogenous tagging of *VPS8A* with FF-ZZ tag

The FF-ZZ tag, containing 3xFLAG (FF), followed by the TEV (Tobacco Etch Virus cysteine protease) cleavage site and the IgG binding domain of protein A (ZZ-domain), was fused to the C-terminus of *VPS8A* macronuclear ORF by homologous recombination, using pVPS8A-ZZflag-Neo4. The construct was obtained by PCR-amplifying the FF-ZZ tag from a donor vector, and subcloning it in the linearized pVPS8A-2mNeon-6myc-Neo4 vector at the MluI and SpeI sites, to replace the 2mNeon-6myc tag. The vector was linearized with SacI and KpnI prior to biolistic transformation of CU428.1. The primers are listed in Supplemental Table S5.

### Expression of Vps16ap-HA and Vps16bp-HA

Vps16ap-HA and Vps16bp-HA were integrated at the metallothionein 1 (*MTT1*) genomic locus[91] in CU428.1 and Vps8ap-FF-ZZ-expressing cell lines by homologous recombination, using VPS16ap-2HA-ncvb and VPS16bp-2HA-ncvb. The 2xHA tag was amplified from a donor vector and subcloned in the linearized ncvb vector at the PmeI and ApaI sites. The *VPS16A* and *VPS16B* ORFs were amplified from genomic DNA, and cloned in the linearized 2HA-ncvb by In-Fusion cloning (Clontech, Mountain View, CA) at the MluI site. The final constructs were linearized with SfiI and biolistically transformed in *Tetrahymena* strains. The primers are listed in Supplemental Table S5.

### Expression of mCherry-Rab7p, mCherry-Rab22Ap, Igr1p-GFP and Stx7l1p-GFP

mCherry-Rab7p, mCherry-Rab22Ap, Igr1p-GFP and Stx7l1p-GFP were integrated at the metallothionein 1 (*MTT1*) genomic locus in CU428.1 and *Δvps8a* strains by homologous recombination, using 2HA-3mCherry-RAB7-ncvb, 2HA-3mCherry-RAB22A-ncvb, and the previously described IGR1-mEGFP-ncvb [92] and STX7L1-mEGFP-ncvb [48] vectors. mCherry-Rab7p and mCherry-Rab22Ap were also transiently expressed in Vps8ap-mNeon expressing cell lines. In these constructs the transgene expression is under the control of the cadmium-inducible *MTT1* promoter[91]. The 2HA-mCherry tag was PCR-amplified and cloned in the PmeI/NheI sites of the p50 vector. Two additional mCherry tags were added at the NheI site of the linearized 2HA-mCherry-p50 by compatible cohesive ends restriction cloning. The 2HA3mCherry tag was then removed from the vector backbone by PmeI/NheI double digestion. The tag was joint to the N-terminus of either *RAB7* or *RAB22A* Macronuclear ORFs by multiple fragment ligation. *RAB7* or *RAB22A* Macronuclear ORFs were previously PCR-amplified and NheI/ApaI digested, and introduced to the Pmel/Apal-digested ncvb vector [46]. The 2HA-3mCherry-RAB7-ncvb,2HA-3mCherry-RAB22A-ncvb, IGR1-mEGFP-ncvb and STX7L1-mEGFP-ncvb constructs were linearized with SfiI and biolistically transformed into *Tetrahymena* strains. The primers are listed in Supplemental Table S5.

### Biolistic transformation

*Tetrahymena* transformants were generated and selected after biolistic transformation as previously described[48, 93, 94]. Transformants were serially transferred 6x/week in increasing concentrations of drug and decreasing concentrations of CdCl_2_ (up to 1.2 mg/ml of paromomycin and 0.1 μg/ml CdCl_2_; up to 18-20 μg/ml of cycloheximide and 0.5-1 μg/ml CdCl_2_; up to 75-85 μg/ml of blasticidin and 0.1 μg/ml CdCl_2_) for at least 3-5 weeks before further testing. At least three independent transformants were tested for each line.

### Fluorescence and Immunofluorescence microscopy

Cells (1.5x10^5^) were washed, fixed with 4% paraformaldehyde for 30 min, and immunolabeled as previously described [47, 52]. Grl3p and Grt1p were visualized using mouse mAbs (mAb) 5E9 (1:10) and 4D11 (1:4), respectively, followed by Texas Red-conjugated goat anti-mouse antibody (1:100) (Life Technologies, Carlsbad, CA). The simultaneous localization of Grt1p and Grl3p was performed as described earlier [47], by directly conjugating purified mAbs 4D11 and 5E9 to Dylight 488 and 650 (Thermo Scientific, Rockford, IL), respectively, and mixing them 1:1 before incubation with samples. For the simultaneous localization of Grl3p with either Igr1p-GFP or Stx7l1p-GFP, the expression of Igr1p-GFP and Stx7l1p-GFP was induced by incubating cells with CdCl_2_ prior to the fixation. Samples were subsequently immunolabeled with 5E9 and Texas Red-conjugated anti-mouse IgG. Specifically, Igr1p-GFP-expressing cells were treated with 1 μg/ml of CdCl_2_ in SPP for 2h at 30°C, and then an additional 1μg/ml of CdCl_2_ was added for 2h. The expression of Stx7l1p-GFP was induced with 1μg/ml of CdCl_2_ for 3h at 30°C. For the simultaneous localization of Grl3p and Cth3p-GFP, cells were stained with 5E9 mAb and with rabbit polyclonal anti-GFP antibody (1:400) (Invitrogen), followed by incubation with Texas Red-conjugated goat anti-mouse and Alexa 488-conjugated donkey anti-rabbit antibodies (1:250) (Invitrogen), respectively, in 1% BSA blocking solution. The expression of mCherry-Rab7p or mCherry-Rab22Ap in wildtype, *Δvps8a*, and Vps8ap-mNeon-expressing cells, was induced with 1μg/ml of CdCl_2_ for 2.5h at 30°C prior to fixation. The visualization of Igr1p-GFP, Stx7l1p-GFP, Sor4p-GFP, Vps8ap-mNeon, mCherry-Rab7p and mCherry-Rab22Ap in fixed cells was not enhanced with immunolabeling. Immunostained cells were mounted with Trolox (1:1000) to inhibit bleaching and imaged on a Zeiss LSM 880 Confocal Laser Scanning Microscope, 100X oil with NA=1.4, with Zen2.1 software (Zeiss, Thornwood, NY). Fixed wildtype, *Δvps8a*, and Vps8ap-mNeon-expressing cells expressing, either mCherry-Rab7p or mCherry-Rab22Ap, were imaged on a Marianas Yokogawa type spinning disk inverted confocal microscope, 100X oil with NA=1.45 with Slidebook6 software (Zeiss, Intelligent Imaging Innovations, Denver, CO). Z-stack images (11-15 stacks along the z-axis at 0.5 μm intervals for Rabs/Vps8ap co-localization analysis; 23-44 and 18-40 stacks along the z-axis at 0.25 μm intervals for Rab22Ap and Rab7p particles analysis, respectively) were processed with Huygens Professional software (Scientific Volume Imaging), colored, denoised and adjusted in brightness/contrast with the program Fiji [95]. Images shown are single slices for clarity.

### Live cell imaging

*Tetrahymena* cells expressing Grl3p-GFP, Sor4p-GFP, Igr1p-GFP, mCherry-Rab7p, or mCherry-Rab22Ap were grown overnight to 1-2x10^5^cells/ml and transferred to S medium for 2h unless otherwise indicated, prior to imaging. Igr1p-GFP expression was induced with CdCl_2_ at 30°C for 4h, as described earlier, while mCherry-Rab7p and mCherry-Rab22Ap expression was induced with 1 μg/ml of CdCl_2_ for 2.5 h. Cells were immobilized in a thin 3% low melting agarose gel pad as described previously [48]. Z-stack images (5-10 stacks along the z-axis at 0.5-2 μm intervals) and time-lapse videos of cells expressing mCherry-Rab7p, and mCherry-Rab22Ap (20 frames, 0.1ms interval), were collected with a Zeiss LSM 880 Confocal Laser Scanning Microsocpe, 100X oil with NA=1.4, with Zen2.1 software (Zeiss, Thornwood, NY). Images and movies were processed with Huygens Professional software (Scientific Volume Imaging), colored, denoised and adjusted in brightness/contrast using Fiji[95]. Images shown are single slices/frames for clarity.

### Labeling endocytic and acidic compartments with FM4-64 and LysoTracker

*Tetrahymena* cells were incubated for 5 min with 5 μM FM4-64, or 7 min with 500 nM LysoTracker (Invitrogen). Cells were washed twice with S-medium, immobilized for live microscopy in 3% low melting agarose gel pads, and imaged within 30 min.

### Electron microscopy

Cells were grown, starved and prepared for electron microscopy as previously described [48]. Fixation was performed at room temperature with 2.5% glutaraldehyde, 1.5% sucrose, and 2% osmium in 6.7 mM sodium cacodylate buffer.

### CLEM imaging

Cells expressing Igr1p-GFP were processed for correlative light and electron microscopy (CLEM) as previously described [48, 96] with a minor modification. Briefly, *Tetrahymena* cells (CU428 or SB281) expressing Igr1-GFP were fixed with 2.5% glutaraldehyde for 1h at room temperature and washed 4x with 0.1 M phosphate buffer, pH 7.4, four times, and then incubated with 0.5µg/ml 4’,6-diamidine-2’-phenylindole (DAPI), a specific DNA-staining fluorescence dye, to stain DNA for 10 min at room temperature. After washing with PB once, the cells were embedded in an agarose gel on a glass bottom dish (MatTek, Ashland, MA, USA). 3D images (50–60 z-stacks × 0.2µm intervals) were obtained using an oil immersion objective lens (PlanApoN60xOSC/NA1.4; Olympus, Tokyo, Japan) and processed by deconvolution. EM observation of the same cells was carried out as described previously [96, 97]. Briefly, the cells in the agarose gel were post-fixed with 1% OsO_4_ (Nisshin EM, Tokyo, Japan) for 1h. The samples were stained en bloc with 2% uranyl acetate (Wako, Osaka, Japan) for 1h, and dehydrated with sequential concentrations of ethanol. The epoxy block containing the cells of interest was trimmed according to the address on the grid and sliced to 80-nm sections. Image data were collected by JEM-1400 (JEOL, Tokyo, Japan) with an accelerating voltage of 80 kV.

To make a correlation between FM and EM images, the aspect ratio of the fluorescence images corresponding to the EM images was adjusted to 1.00 to 1.23 or to 1.13 for Fig. 2I and 2J, respectively, using Powerpoint. Then, a display mode of FM images was inverted from the negative to the positive one using Powerpoint. Finally, the FM images were overlaid to the EM images using Powerpoint with 50% transparency to generate a single montage image.

### Dibucaine assay

Stimulation of mucocyst exocytosis with dibucaine was performed as previously described[47].

### Immunoprecipitation

The following protocol was used in immunoprecipitation of Vps8ap-GFP, Sor4p-GFP, Vps8ap-mNeon and Sor4p-mNeon, prior to visualization by Western blotting. Cells were grown overnight to 1.5-2x10^5^ cells/ml. Vps8ap-GFP and Sor4p-GFP were immunoprecipitated with GFP-nAb agarose Spin Kit (Allele Biotechnology Inc., San Diego, CA) from detergent cell lysates, as follows. 400-ml cultures were starved for 1 h in 10 mM Tris, pH 7.4, pelleted and resuspended in lysis buffer [20 mM Tris-HCl pH 7.4, 50 mM NaCl, 1 mM MgCl_2_, 1 mM DTT, 1 mM EGTA, 0.2% NP-40, 10% glycerol], supplemented with cOmplete EDTA-free protease inhibitor cocktail tablets (Roche). After 45 min at 4°C, lysates were cleared by centrifugation at 30000 rpm (Beckman type 45 Ti rotor) for 1 h. 80 μl of pre-washed GFP-nAb agarose resin were added to the cleared lysates and mixed for 3h at 4°C. The beads were then washed once with wash buffer (10 mM Tris-HCl pH 7.4, 500 mM NaCl), transferred to a spin column and washed five times with wash buffer. GFP-tagged proteins were eluted by adding 500 μl of 0.2 M glycine, pH 2.5. Proteins were precipitated with 10% trichoroacetic acid (TCA) and 0.02% deoxycholate (DOC), and resuspended in 40 μl 100°C SDS-PAGE sample buffer.

Vps8a-mNeon and Sor4p-mNeon were immunoprecipitated in parallel from cell lysates with nProtein A Sepharose 4 Fast Flow resin (GE Healthcare Life Sciences). Sor4p-mNeon expression was induced with 0.5μg/ml CdCl_2_ for 3h. Cells expressing Vps8ap-mNeon from the *VPS8A* locus were similarly treated with CdCl_2_, to avoid differences due to cadmium stress. 200 ml cultures were washed once with 10mM Tris, pH 7.4 and lysed as described earlier. The cleared lysates were mixed with 50 μl nProtein A Sepharose beads, which were pre-conjugated with ~50 μg mouse anti-c-myc mAb (9E10) for 2h at 4°C, according to the manufacturer’s instructions. The beads were then washed three times with 20 mM Tris-HCl pH8, 1 mM EDTA, 50 mM NaCl, 0.1% NP-40, 1 mM DTT, 10% glycerol supplemented with cOmplete EDTA-free protease inhibitor cocktail tablets (Roche). Proteins were eluted with 400μl 0.1M glycine pH 2.5, precipitated with 10% TCA and 0.02% deoxycholate, and dissolved in 40μl 100°C SDS-PAGE sample buffer.

### Co-immunoprecipitation (Co-IP)

Vps8ap-FF-ZZ fusion protein was immunoprecipitated from detergent lysates of cells that were also transiently expressing either Vps16ap-HA or Vps16bp-HA, using anti-FLAG beads (EZ view Red Anti-FLAG M2 Affinity Gel, Sigma). In brief, 250ml cultures were grown overnight to 1x10^5^ cells/ml, then treated with 1 μg/ml of CdCl_2_ for 2.5h. They were washed once with 10 mM Tris, pH 7.4, pelleted and resuspended for 45min in cold lysis buffer (20 mM Tris-HCl pH 7.4, 50 mM NaCl, 1 mM MgCl_2_, 1 mM DTT, 1 mM EGTA, 0.2% NP-40, 10% glycerol, 4% BSA), supplemented with protease inhibitor cocktail tablets (Roche). Lysates were cleared by centrifugation at 35000 rpm (Beckman type 45 Ti rotor) for 1.5 h. 75 μl of anti-FLAG beads, pre-incubated with cold lysis buffer for 2h, were added to the cleared lysates and mixed for 2h at 4°C. The beads were then washed five times with 20 mM Tris-HCl pH8, 1 mM EDTA, 1 M NaCl, 0.1% NP-40, 1 mM DTT, 10% glycerol, and resuspended in 100 μl 100°C SDS-PAGE sample buffer.

### Western blotting

For Western blotting of whole cell lysates, 5x10^5^ cells from overnight cultures were washed once with 10 mM Tris pH 7.4, resuspended in 500 μl 10 mM Tris, and precipitated with 10% TCA on ice for 30 min. Samples were then centrifuged at 12000 rpm (Eppendorf F-45-30-11 rotor) in an Eppendorf microfuge for 10 min at 4°C, washed once with ice-cold acetone and resuspended in 100 μl 100°C SDS-PAGE sample buffer. Proteins were resolved with the Novex NuPAGE Gel system (4-20% Tris-Glycine gels, Invitrogen), and transferred to 0.2 μm PVDF membranes (Thermo Scientific). Blots were blocked with 5% dried milk or 3% BSA in 1X TBS-Tween (25 mM Tris, 3 mM KCl, 140 mM NaCl, 0.05% Tween 20, pH 7.4). The rabbit anti-Grl1p serum, the mouse mAb anti-GFP (BioLegend), the mouse mAb anti-c-myc (9E10, Sigma), the rabbit anti-FLAG (Sigma), and the mouse mAb anti-HA (HA.11, BioLegend) antibodies, were diluted 1:2000, 1:5000, 1:1000, 1:2000, 1:2000, respectively, in blocking solution. Proteins were visualized with either anti-rabbit IgG (whole molecule)-HPeroxidase (Sigma) or ECL Horseradish Peroxidase-linked anti-mouse (NA931) (GE Healthcare Life Sciences, Little Chalfont, UK) secondary antibody diluted 1:20000, and SuperSignal West Femto Maximum Sensitivity Substrate (Thermo Scientific).

### Colocalization analysis

To estimate the extent of colocalization, the Fiji-JACoP plugin was used to calculate Manders’ coefficients M1 or M2 [98]. M1 is defined as the ratio of the summed intensities of pixels from the green image for which the intensity in the red channel is above zero to the total intensity in the green channel, and M2 is defined conversely for red. The images were corrected for noise, and the M1 and M2 coefficients were calculated setting the threshold to the estimated value of background. Between 25 and 65 non-overlapping images were analyzed for each sample.

### Particle analysis

The estimation of the number and size of particles was obtained by using the Fiji tool “Analyze Particles” (<u>http://imagej.nih.gov/ij/</u>). The number of particles was calculated using non-overlapping images, which were corrected for noise and analyzed by setting the threshold to the estimated background value, and then converted to “Mask”. The calculation was restricted to different area-based size ranges, selected on the base of the overall population size in both WT and *Δvps8a* cells, including particles between 0.1 and 3 μm^2^ for Sor4p-GFP particles, between 0.2 and 3 μm^2^ for mCherry-Rab22Ap particles, and between 0.2 and 8 μm^2^ for mCherry-Rab7p-particles.

The number of particles with a given size is reported as Mean value.

### Homology searching and Phylogenetic tree construction

For the comparative genomic detection of VPS8 homologues in the Oligohymenophorea, Vps8 sequences from diverse eukaryotes were used as queries against translated ORF coding sequences from the genomes and transcriptomes of selected ciliate species using the BLAST algorithm. Positive BLAST hits against a Vps8 query were those with an E value of less than 0.05, for which reciprocal BLAST against the genome containing the query sequence retrieved either the same sequence or an isoform of the sequence with the same E value or lower. To identify a hit as orthologous to the query, we further required that the E value be at least three orders of magnitude lower than the next lowest hit. Hits that were consistent with the first two criteria but did not show clear superiority over other hits were classified as potential hits, which may be either Vps8 or 41, or Vps3 or 39.

Once homologues of HOPS/CORVET had been determined via BLAST, the phylogenetic relationships of the VPS8 homologues within the Oligohymenophorea were determined using both maximum likelihood (via the RAxML algorithm) and Bayesian inference (via the MrBayes algorithm) run using the CIPRES server. Homologues were aligned using MUSCLE and alignments were manually trimmed to retain regions of unambiguous homology. For the RAxML trees, ProtTest was used to select an appropriate rate evolution model. Consensus trees were constructed through manual inspection of both RAxML and MrBayes outputs and determination of corresponding nodes, mapped onto the MRBayes topology. The root of the tree was determined using *Oxytricha trifallax* Vps8 sequences as an outgroup, showing that expansions within the Spirotrichea were lineage specific and the post-Spirotrichea expansions form a single clade with MrBayes/RAxML support values of 1/100.

## Author Contributions

DS, HO, MI, GRB, and CK designed and conducted experiments; MI, TH and APT designed experiments; ER, XL, JKP, and JBD designed computational approaches; ER and XL conducted computational approaches; WAB and DHL provided genomic resources; DS, ER, HO, and MI prepared figures; SB, DS, ER, XL, MI, TH, JBD and APT wrote the paper.

## Acknowledgements

DS and APT would like to thank Harsimran Kaur for the RAB22A and RAB7 constructs, Lev Tsypin and Harsimran Kaur for valuable discussion, Vytas Bindokas and Christine Labno, as well as Yimei Chen, for assistance with light and electron microscopy, respectively, at University of Chicago Core Facilities; K. Mochizuki (Institute of Human Genetics, CNRS-University of Montpellier, France) for NEO4 plasmids, and J. Frankel (University of Iowa, Iowa City) for hybridomas 5E9 and 4D11. ER and JBD thank Michael Lynch (Arizona State Univ.) for access to *P. caudatum* genomes. Work in TH’s laboratory was supported by JSPS Kakenhi Grant Numbers, JP24570227, JP15K07066 to MI, and JP26291007, JP25116006, JP15K21730 to TH. ER was supported by an Alberta Innovates - Technology Futures Graduate Student Scholarship and a Vanier Canada Graduate Scholarship. work in J.K.P.’s laboratory was supported by the Howard Hughes Medical Foundation and NIH 1RO1ES025009. Work in JBD’s laboratory was supported by a Discovery Grant from the Natural Sciences and Engineering Research Council of Canada (RES0021028) and JBD is the Canada Research Chair (Tier II) in Evolutionary Cell Biology. Work in APT’s laboratory was supported by NIH 1RO1GM105783.

